# Polar protein Wag31 both activates and inhibits cell wall metabolism at the poles and septum

**DOI:** 10.1101/2022.05.31.494259

**Authors:** Neda Habibi Arejan, Parthvi Bharatkumar Patel, Samantha Y. Quintanilla, Arash Emami Saleh, Cara C. Boutte

## Abstract

Mycobacterial cell elongation occurs at the cell poles; however, it is not clear how cell wall insertion is restricted to the pole and organized. Wag31 is a pole-localized cytoplasmic protein that is essential for polar growth, but its molecular function has not been described. In this study we used alanine scanning mutagenesis to identify Wag31 residues involved in cell morphogenesis. Our data show that Wag31 has separate functions in not only new and old pole elongation, but also inhibition of both septation and new pole elongation. Our examination of phospho-ablative and phospho-mimetic mutants of Wag31 suggests that phosphorylation of Wag31 promotes old pole elongation, while the unphosphorylated form of Wag31 may promote resolution of the septum. This work establishes new regulatory functions of Wag31 in the mycobacterial cell cycle and clarifies the role of phosphorylation on Wag1.

**Importance:** Despite many previous studies, the molecular mechanisms of polar growth in mycobacteria is unclear. Wag31 is required for this polar elongation. In this work, we dissect Wag1 function by phenotyping *wag31* point mutants. We find that Wag31 promotes elongation at both poles in different ways, and it can also inhibit cell wall metabolism at both the new pole and the septum. This work is important because it clarifies that Wag31 is doing several different things in the cell and gives us genetic tools to disentangle its functions.

## Introduction

To expand in size bacteria need to cut and add to the existing peptidoglycan cell wall without disturbing its integrity. Peptidoglycan is critical for maintenance of cell shape, and is made of sugar chains crosslinked by small peptides. Many rod shaped organisms perform this cell wall expansion all along the lateral walls (1, 2). Mycobacteria, as well as many other Actinomycetes and some Alphaproteobacteria, restrict cell wall expansion to the cell poles. In mycobacteria, the old pole elongates continuously, but there is a delay in initiation of elongation at the new pole – the one most recently created by cell division (3) - which results in asymmetric cells (3–5). The molecular mechanisms that control polar cell wall expansion are not well described. In this paper we dissect the function of Wag31 (AKA DivIVA, Rv2145c, MSMEG_4217), which is essential for polar growth in mycobacteria (6, 7), though its molecular function is unknown. Wag31 a mycobacterial homolog of DivIVA. DivIVA proteins are found in Firmicutes and Actinobacteria (Fig. S1). They typically localize to the poles and/or septum, and function to recruit and regulate other factors that have functions in cell division, chromosome segregation and cell wall synthesis (8–12).

Wag31, is localized at the poles (13, 14) and at the septum just before division(15). In addition to its role in polar growth, it also helps with chromosome segregation by recruiting the segregation factor ParA to the poles (16). Wag31 is not required for cell wall metabolism, but it is needed to direct this metabolism to the pole (17), though it is not known how it does this. *Mycobacterium smegmatis* (*Msmeg*) does not regulate polar cell wall synthesis by restricting its essential peptidoglycan transglycosylases to the poles, as some other pole-growing species do (9, 12, 18, 19), since both RodA and PBP1 are delocalized around the entire cell instead of restricted to the poles (17). However, *Msmeg* does restrict synthesis of many cell wall precursor enzymes near the poles, through association with the Intracellular Membrane Domain (IMD), a chemically distinct region of the plasma membrane found at the sub polar regions in growing mycobacterial cells (17, 20). Depletion of Wag31 destabilizes the IMD (21), though it is not clear whether Wag31 regulates this membrane domain directly.

Polar cell wall elongation is highly regulated and is known to decrease under stress (17, 22). Phosphorylation of Wag31 at T73 (23) has been proposed as a possible means of controlling polar elongation (24). In other species, DivIVA homologs involved in cell morphogenesis have been shown to be regulated by phosphorylation (23, 25–31). Here, we identify a series of point mutants of *wag31* that exhibit diverse phenotypes, and indicate that Wag31 has roles in inhibition of new pole elongation and septal inhibition, in addition to its established roles in promoting elongation at the poles. Also, we show that while phosphorylation of Wag31 does moderately affect polar growth, it is not a critical upstream regulator of this process.

## Results

### Wag31 has separate roles in several steps of the cell cycle

Depletion of Wag31 leads to arrest of polar elongation, loss of pole structure, (6, 24) and delocalization of peptidoglycan and mycomembrane metabolism (7). To dissect Wag31’s functional roles in these processes, we profiled the phenotypes of *Msmeg* strains with *wag31* WT replaced by the alanine mutants of *wag31*, using L5 allele swapping (32) (Fig. 1A). We tested the stability of the mutated Wag31 proteins and only characterized mutants with stable Wag31 (Fig. S2). We were unable to swap some of the mutants, thus P2, T4, K15, and EQR 210-3 might be essential for Wag31 function (Fig S3C).

**Figure 1.**
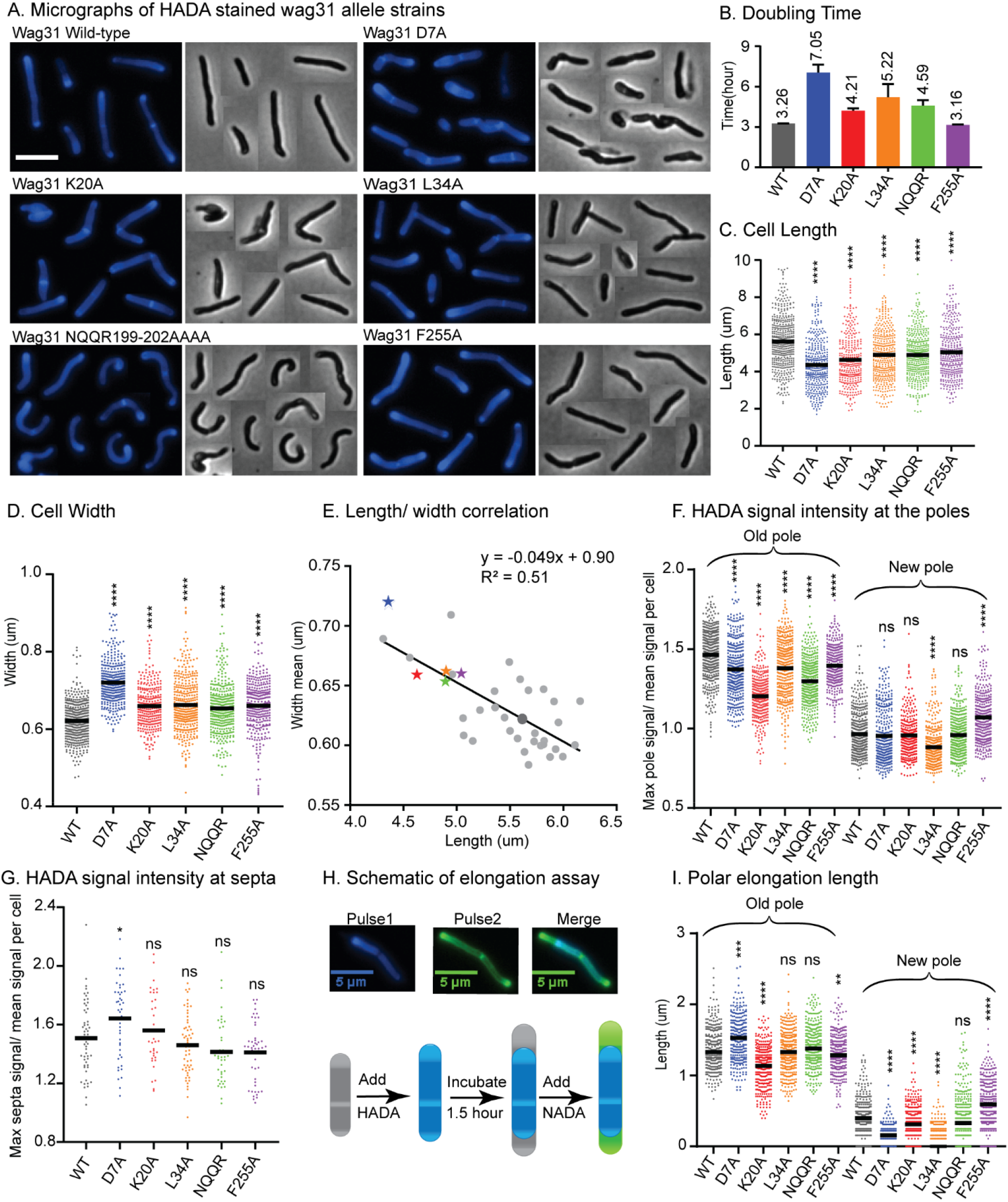
Wag31 has multiple distinct roles in both elongation and division. (A) Phase (right) and fluorescence (left) images of *Msmeg wag31* allele strains stained with HADA. The scale bar is 5 microns, and it applies to all images. (B) Doubling times of *Msmeg* cells expressing WT or *wag31* alanine mutants. The means (on top of bars) are an average of three biological replicates. Error bars represent SD. (C) Cell lengths of the *wag31* allele strains. Black bars are at the mean. (D) Cell widths of the *wag31* allele strains. Black bars are at the mean. (E) Correlation between cell length and cell width. Black bar is linear regression fit. The five mutants in the other panels are shown as colored stars, the rest of mutants are shown as light gray dots, *wag31* WT is a dark gray dot. Relative polar (F) and septal (G) HADA intensity of *wag31* allele strains. (H) Schematic of elongation assay dual-FDAA staining method to assay elongation. (I) Length of polar elongation in the *wag31* allele strains, as measured by the elongation assay in H. Black bar is at the median. ns, p >0.05, *, P = < 0.05, **, P=< 0.005, ***, P =<0.0005, ****, P = <0.0001. All *P*-values are calculated by one-way ANOVA, Dunnett’s multiple comparisons test.

We phenotypically profiled the *wag31* allele swap strains (Fig 1, S3, S4, S5). Most of the mutants at the N-terminus of Wag31 impair growth rate, while most mutants in the middle and C-terminus of the protein do not (Fig. 1B, S3A). Microscopy experiments show that many of the mutant cells are short and/ or wide (Fig. 1BC, S3A). Most of the *wag31*_Mtb_ mutant strains have higher signal than the WT after staining with the fluorescent D-alanine HADA (33) (Fig. S4A, S5A, S6, S7), which may indicate de-repressed peptidoglycan metabolism or increased cell wall permeability. To compare localized peptidoglycan between strains, we used relative HADA intensity at the poles and septa: this is the maximum intensity at that region divided by the mean intensity of that cell. Because the older pole almost always stains more brightly by HADA (34); throughout we assume that the brighter pole is the old pole.

Most mutants had lower relative HADA signal than WT at the old pole, and none have higher signal (Fig. 1F, S4B). At the new pole most mutants stain like the WT, but a few near the N-terminus are dimmer than WT, and a few near the C-terminus are brighter than WT (Fig. 1F, S4C). This data indicates that mutants in Wag31 can have different impacts on the peptidoglycan metabolism at the two cell poles. Septal HADA intensity is increased in a few mutants (D7A, K20A, DER 185,7,8), while it is decreased in others (Fig. S5B), suggesting that Wag31 regulates peptidoglycan synthesis at the septa as well. We also observe that the septal placement is shifted toward either pole in some of the mutants (Fig. S5C).

A variety of different phenotypes indicative of assorted functional roles for Wag31 were apparent across all the mutants. We chose five strains with mutations in residues that are highly conserved among mycobacteria species (Fig. S1) and which represented the range of observed phenotypes for further study: D7A, K20A, L34A, NQQR199-202AAAA, and F255A.

### Wag31 is involved in septal inhibition via glutamate 7

The *wag31* D7A strain grows slowly, and the cells are short and wide (Fig. 1ABCD). There is a slight decrease in HADA intensity, compared to WT, at both poles (Fig. 1F), but HADA intensity is significantly increased at the septa (Fig. 1G). To determine whether HADA staining at the poles is indicative of polar elongation or peptidoglycan remodeling (34), we performed an elongation test with two colors of fluorescent D-amino acids. We stained first with HADA, then outgrew the cells without stain for 1.5 hours, then stained with the green fluorescent D-alanine NADA, and measured the length of the poles that are green and not blue (Fig. 1H). Results from this experiment show that the *wag31* D7A strain elongates ~.14 μm more than the WT at the old pole, while elongation at the new pole is ~.27 μm less than the WT (Fig. 1I). Our data show that the net elongation defect (7%) does not have major role in substantially decreased observed cell length (22%) (Fig. 1C), indicating that division must be initiated in shorter cells in this strain. The septal HADA staining suggests that there is more peptidoglycan metabolism at the septum than in WT cells (Fig. 1G, S5B) and that septal staining occurs in shorter cells (Fig. S6). This cell division may sometimes be too early, which could lead to the unusual number of ghosts in the *wag31* D7A phase images (Fig. 1A). These data suggest that Wag31 is involved in inhibition of septation through D7. The effects of this mutation could be due to an inability to interact with other proteins or the cell membrane. We localized Wag31 D7A-GFPmut3 in a merodiploid strain – because the allele swaps were not viable - and find that it localizes like the Wag31 WT-GFPmut3 at the poles and septum (Fig. 4), which allows us to infer that Wag31 D7A is likely defective in protein-protein interactions.

### Wag31 regulates polar elongation through N-terminal residue K20

The N-terminus of Wag31 is highly conserved and it has been structurally characterized (35, 36). In *Bacillus subtilis*, F17 is involved in interaction with the membrane at the cell pole (35). In *Mycobacterium smegmatis*, K20 is in a similar position as F17 in DivIVA_*Bsub*_ in the structure (36).

Our microscopy data show that the *wag31* K20A mutant has decreased HADA staining compared to WT at the old pole, but equivalent staining at the new pole (Fig. 1F). The elongation test shows that the *wag31* K20A mutant elongates ~.26 μm less than the WT at the old pole, and ~.14 μm less at the new pole (Fig. 1FHI). The defect in *wag31* K20A elongation (22% less) is similar to the defect in cell length (18%), indicating that this mutant is defective in elongation, especially at the old pole. The *wag31* K20A strain has similar septal intensity as the WT (Fig 1G), implying that peptidoglycan metabolism is comparable; however, the septal location is shifted slightly toward the old pole (Fig. S5C), likely because the old pole is elongating more slowly.

To determine if K20 is involved in membrane association, we localized K20A-GFPmut3 in a merodiploid strain. Wag31 first associates with the membrane at the septum, when new poles are formed during division (15). Our data show that Wag31 K20A mutant protein is defective in septal localization, whereas its localization at the poles is similar to the Wag31 WT (Fig. 4D). This data indicates that the Wag31 K20A mutant may be slightly impaired in its ability to adhere to the polar membranes. The polar growth phenotypes in the *wag31* K20A mutant could be due to defects in homo- or hetero-oligomeric interaction, or changes in its affinity to the membrane.

### Wag31 regulates new pole elongation through L34

We found that the *wag31* L34A mutant also has a defect in polar elongation. The *wag31* L34A strain has decreased HADA staining at both poles, the old pole staining is 1.5% less, and the new pole is ~11% less than the WT (Fig. 1F). The old pole of the *wag31* L34A strain elongates the same amount as the WT strain, but the new pole has ~76% less elongation (Fig. 3I). In the population of *wag31* L34A mutants that we characterized, ~65% had no observable elongation at the new pole, while only 11% of cells in the *wag31* WT population had no new pole elongation in the 1.5 hour assay. Thus, it appears that the Wag31 L34 residue contributes to both initiation and extension of the new pole. The short cell length and slow growth rate (Fig. 1ABC) are likely due to this defect in new pole elongation, as we did not observe defects in the intensity or location of septa (Fig. 1G, S5C). We localized the Wag31 L34A-GFPmut3 in a merodiploid, and it does not have defects in polar localization. It is slightly dimmer than the Wag31 WT-GFPmut3 at the septum (Fig. 4D), which is the site of new pole construction.

### Wag31 inhibits new pole initiation through F255

We made several alanine mutants in the C-terminus of Wag31, and we found that in the *wag31* F255A mutant, there is increased intensity of HADA at the new pole, and less at the old pole (Fig. 1F). The elongation test shows that the *wag31* F255A strain elongates ~47% more than the WT at the new pole, while the old pole elongates 6% less compared to the WT strain (Fig. 1I). There is no difference in polar and septal localization between the Wag31 F255A-GFPmut3 and the Wag31 WT-GFPmut3 (Fig. 4). The HADA septal intensity is similar in the *wag31* F255A mutant compared to the WT (Fig. 1G), while septal location is shifted toward to the old pole, likely due to the increased elongation activity at the new pole (Fig. S5C). The *wag31* F255A grows as fast as the WT strain (Fig. 1B), and the cells are slightly shorter (10%) and thicker than the WT (Fig. 1CDE). The increased elongation must be accompanied by septation at a shorter cell length to cause the shorter cells seen in this mutant (Fig. 1C). These data indicate that Wag31 has inhibitory role on elongation of the new pole through its C-terminal residues (Fig. S4). So, Wag31 can both promote and inhibit polar growth.

### The formation of rod-shape morphology is affected by NQQR199-202 residues

Wag31 is necessary for rod shape (6, 7, 13). Forming a straight rod requires that new cell wall material be added evenly around the circumference of the cell. Previous work showed cell bending when Wag31-eGFP replaces native Wag31 (14), suggesting that deforming the Wag31 network may lead to uneven cell wall insertion. We observed a similar curved cell morphology in the *wag31* NQQR199-202AAAA mutant (Fig. 1A). While the *wag31* NQQR199-202AAAA mutant has slightly decreased HADA intensity at the old pole (Fig. 1F), our elongation test assay shows that elongation is similar to the Wag31 WT at both poles (Fig. 1I). There are no differences in localization of Wag31 NQQR199-202AAAA-GFPmut3 and Wag31 WT-GFPmut3 (Fig. 4). We propose that the Wag31 NQQR199-202AAAA mutation may cause cell wall insertion to be uneven around the circumference of the cell, leading to slight polar bulging and curved cells (Fig. 1A).

### Phospho-site T73 of Wag31 affects both polar and septal peptidoglycan metabolism

Wag31 is known to be phosphorylated by the Serine Threonine Protein Kinase PknA at T73 (23). In previous work in *Mycobacterium smegmatis*, the WT *wag31* was transcriptionally depleted while *wag31* T73A (phospho-ablative) or *wag31* T73E (phospho-mimetic) were induced, and peptidoglycan metabolism was assessed by BODIPY-Vancomycin staining. This work suggested that Wag31~P promoted, and unphosphorylated Wag31 inhibited, polar peptidoglycan synthesis (24). However, that study did not control for the amount of different versions of Wag31 present in the cell at the time of staining. We built the *wag31* T73A and *wag31* T73E strains through allele swapping with the native promoter to normalize the amount of protein and found they have similar growth rates to the WT (Fig. 2AC), but both mutants are slightly longer (Fig. 2D). We observed that the *wag31* T73E strain has slightly greater HADA staining at the old pole, while both strains have similar signal to the WT at the new pole (Fig. 1G). The elongation test showed that the *wag31* T73E strain elongates ~9% more and the wag31 T73A strain elongates ~8% less than the WT at the old pole. We did not observe significant differences in elongation at the new pole (Fig. 2H). The increased cell length (Fig. 2D) and slightly decreased elongation of the wag31 T73A strain (Fig. 2H) suggest that this mutant also activates division at longer cell lengths.

**Figure 2.**
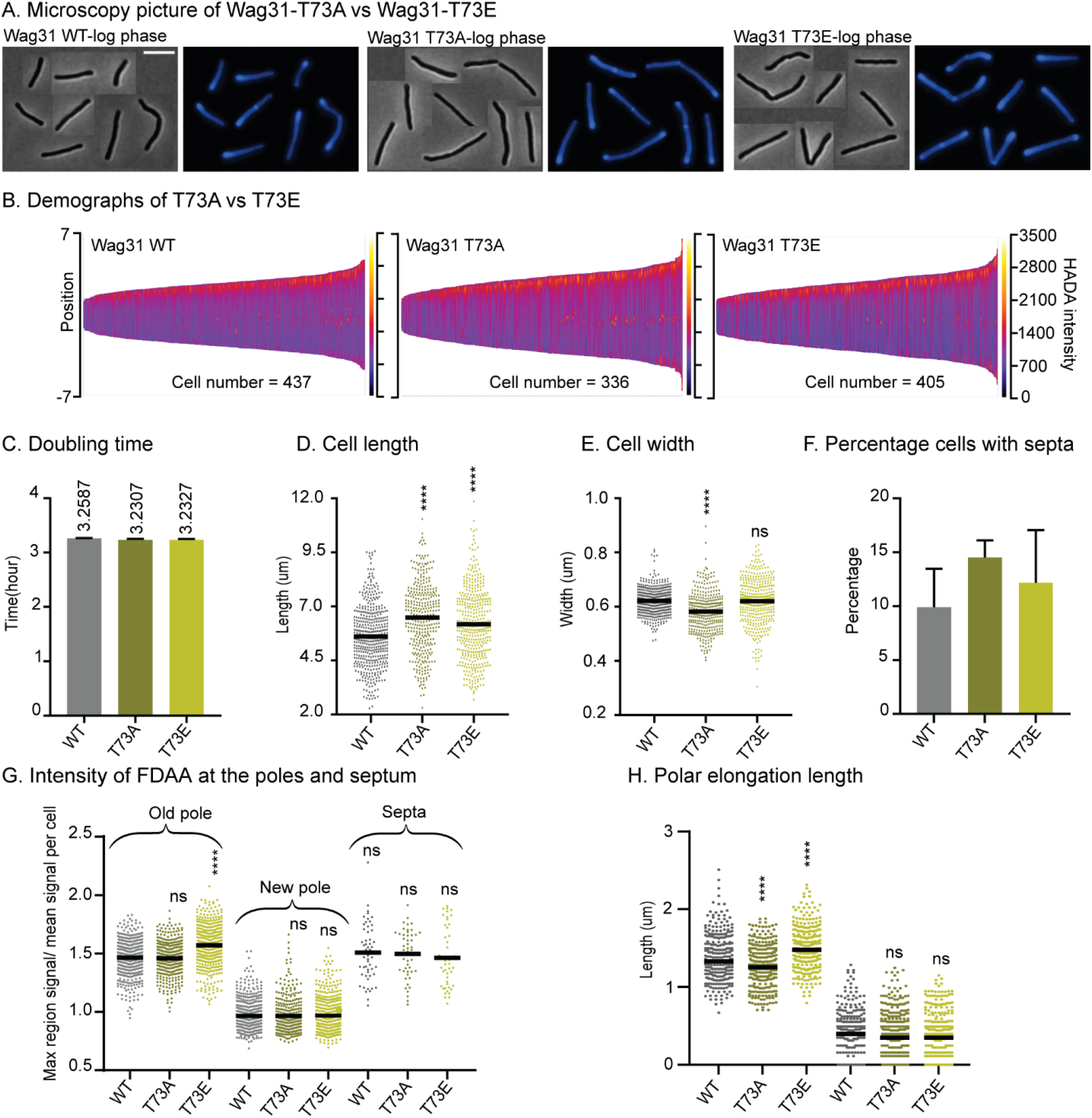
Phosphorylation of T73 on Wag31 subtly affects the cell cycle, but does not affect growth rate. (A) Phase (right) and fluorescence (left) images of *Msmeg wag31* allele strains stained with HADA. The scale bar is 5 microns, and it applies to all images. (B) Demographs of HADA intensity (color scale) across the length of the cell (Y axis) of the *wag31* allele strains. The cells were sorted by length, with shortest cells on the left and longest on the right of each demograph. Cells were also pole-sorted according to HADA intensity, such that the brighter pole (presumed to be the old pole) was oriented to the top along the Y axis. At least 100 cells were analyzed from each of three independent biological replicates of each strain. (C) Doubling times of *Msmeg* cells expressing WT or *wag31* mutants. The means (on top of bars) are an average of three biological replicates. Error bars represent SD. (D) Cell lengths of the *wag31* allele strains. Black bars are at the mean. (E) Cell widths of the *wag31* allele strains. Black bars are at the mean. (F) Percentage of cells in (G) that have septal HADA signal. (G) Relative polar and septal HADA intensity of Wag31 WT, Wag31 T73A, and Wag31 T73E. (H) Length of polar elongation in the *wag31* allele strains, as measured by the pulse-chase-pulse method. Black bar is at the median. ns, p >0.05, ****, P = <0.0001. All *P*-values are calculated by one-way ANOVA, Dunnett’s multiple comparisons test.

In order to determine whether phosphorylation of Wag31 might regulate cell wall metabolism under stress, we performed HADA staining and microscopy of the same strains in stationary phase (Fig. 3A). The results show that these phospho-mutants of *wag31* have the same cell length as WT (Fig. 3C), indicating that phosphorylation on Wag31 is not important for regulating cell length and polar growth in stationary phase. We found slight differences in HADA staining at the old pole between the mutants and the WT, which are curiously in the opposite direction as in log. phase, with the *wag31* T73A phospho-ablative mutant staining more brightly and the phospho-mimetic mutant staining more dimly (Fig. 3F). This suggests that phosphorylation on Wag31 may have some minor role in regulating cell wall metabolism in stress, but that it is likely downstream of several other regulators. The *wag31* T73A mutant has a higher percentage of cells with active septa in stationary phase (Fig. 3EF), indicating that the function of unphosphorylated Wag31 in slowing septation may continue into stationary phase, but that this regulation is not sufficient to affect cell length.

**Figure 3.**
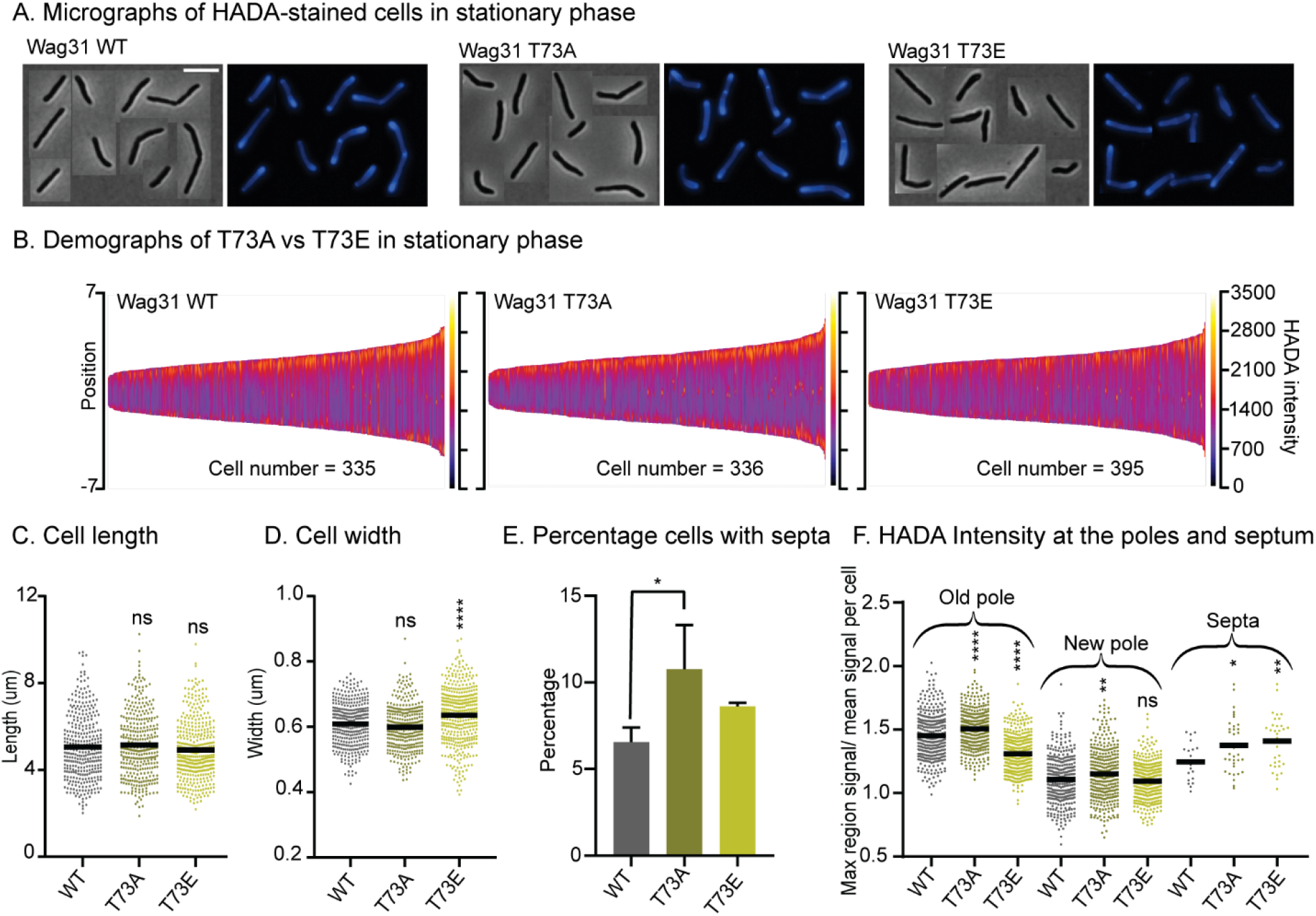
Wag31 phospho-site T73 is not an upstream regulator of peptidoglycan metabolism in stationary phase. (A) Phase (right) and fluorescence (left) micrographs of *Msmeg* cells in stationary phase which are expressing Wag31 WT, Wag31 T73A, and Wag31 T73E. (B) Demographs of HADA intensity (color scale) across the length of the cell (Y axis) of the *wag31* allele strains. The cells were sorted by length, with shortest cells on the left and longest on the right of each demograph. Cells were also pole-sorted according to HADA intensity, such that the brighter pole (presumed to be the old pole) was oriented to the top along the Y axis. At least 100 cells were analyzed from each of three independent biological replicates of each strain. (C) Cell lengths of the *wag31* allele strains. Black bars are at the mean. (D) Cell widths of the *wag31* allele strains. Black bars are at the mean. (E) Percentage of cells in (G) that have septal HADA signal. (F) Relative polar and septal HADA intensity of Wag31 WT, Wag31 T73A, and Wag31 T73E. ns, p >0.05, *, P = < 0.05, **, P=< 0.005, ****, P = <0.0001. All *P*-values are calculated by one-way ANOVA, Dunnett’s multiple comparisons test.

**Figure 4.**
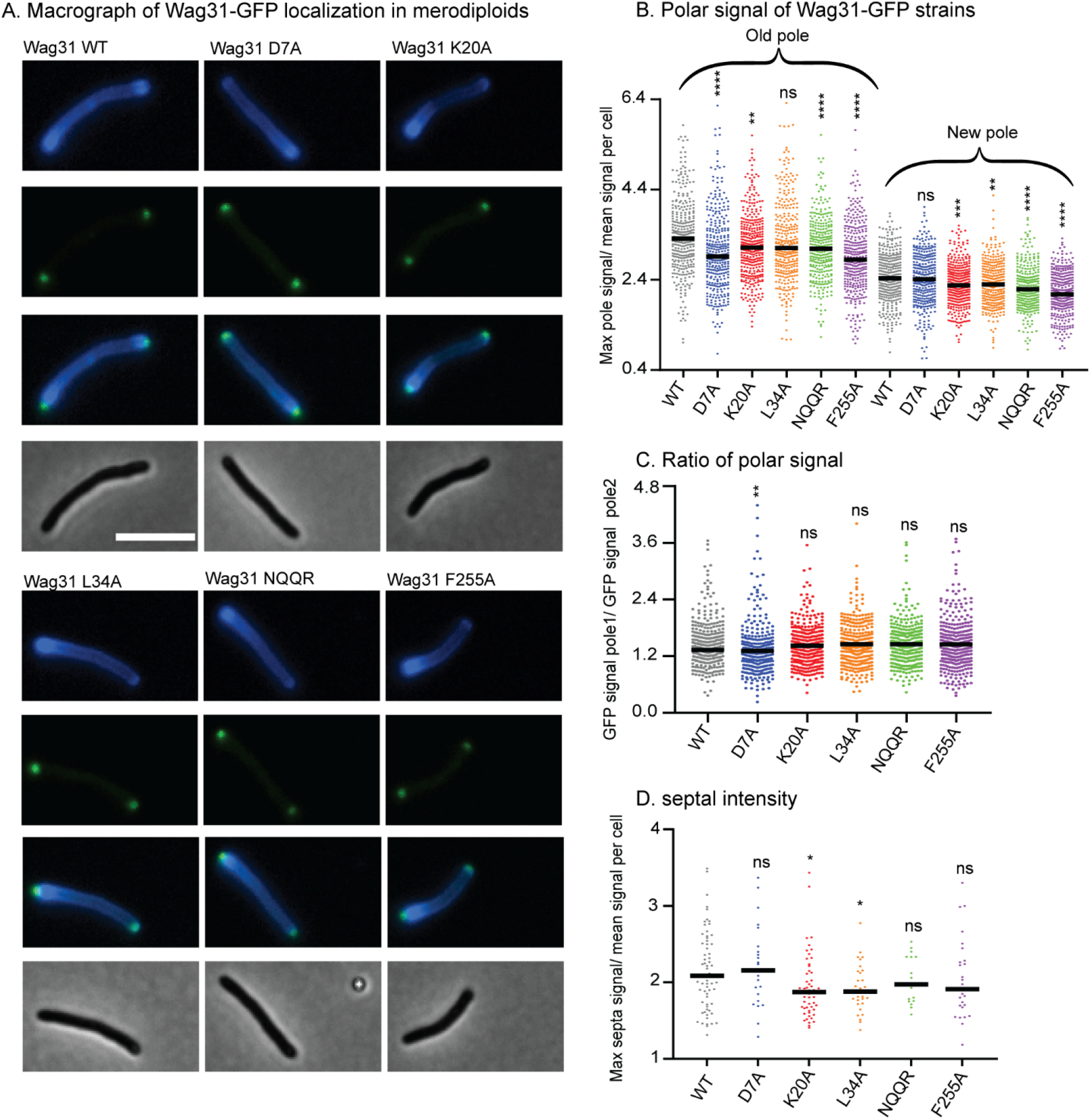
Wag31 mutant proteins localize normally in merodiploid strains. (A) Micrographs of WT *Msmeg* expressing Wag31-GFPmut3 constructs with the indicated mutations. Top: HADA; second: GFP; third: merged; bottom: phase. The scale bar is 5 microns, and it applies to all images. (B) Relative polar intensity of GFPmut3 signal from cells in A. (C) Ratio of normalized signal at the old pole over normalized signal at the new pole, from B. (D) Relative septal intensity of GFPmut3 signal from cells in A. ns, P >0.05, *, P =< 0.05, **, P=< 0.005, ***, P=< 0.0005, ****, P= <0.0001. All *P*-values (B), (C) are calculated by one-way ANOVA, Dunnett’s multiple comparisons test and the *P*-value (D) is calculated by the Welch’s t-test.

Recent work suggests that Wag31 is a favored substrate of the Serine Threonine Phosphatase PstP which localizes to the septum during late division (37). This suggests that Wag31 is dephosphorylated at the septum and may primarily be phosphorylated at the poles. It may be that the phosphorylation only accumulates sufficiently to affect peptidoglycan metabolism once the pole has matured into an old pole, since there are no differences in HADA staining between the WT and phospho-mutants at the new pole (Fig. 2GH). In summation, our data with the phospho-mutants indicates the phosphorylation on T73 of Wag31 modestly promotes polar growth at the old pole, while the unphosphorylated form may slightly inhibit both polar elongation and septation.

### Wag31’s effects on MurG and GlfT2 localization do not correlate with effects on polar growth

Recent work suggests that localization of the enzymes that synthesize cell wall precursors to the subpolar Intracellular Membrane Domain (IMD) may be critical in restricting elongation to the pole (17, 20). Depletion of Wag31 has been shown to delocalize the IMD(21), though it is not known if Wag31 regulates the IMD, or if the IMD localization is merely dependent on an intact cell pole.

To probe whether Wag31 regulates IMD structure, we localized both GlfT2-mcherry2B and MurG-Venus in selected Wag31 mutants. GlfT2 is the last cytoplasmic galactan enzyme in arabinogalactan synthesis (38), associates with the IMD, and localizes in the typical pattern of IMD-associated proteins (20) in the subpolar region and spottily along the lateral walls. MurG is the final cytoplasmic enzyme in peptidoglycan precursor synthesis, has a similar localization pattern as GlfT2, (14) and is found in both cytoplasmic and IMD membrane fractions (21).

Our microscopy data show that localization of GlfT2-mcherry2B is significantly decreased at the old pole in the Wag31 K20A, D7A, L34A and F255A mutants compared to the WT, while it was unchanged in the rest of the mutants at the new pole (Fig 5AE). Localization of MurG-Venus is slightly decreased at the old pole in the *wag31* D7A, L34A and NQQR199-202 mutants compared to the WT (Fig. 5C). At the new pole, localization of GlfT2-mcherry2B is largely unaffected at the new pole, with modest defects in localization in the *wag31* D7A, T73A, and T73E strains (Fig. 5BE). Localization of MurG-Venus is unchanged in all mutants at the new pole (Fig. 5D).

**Figure 5.**
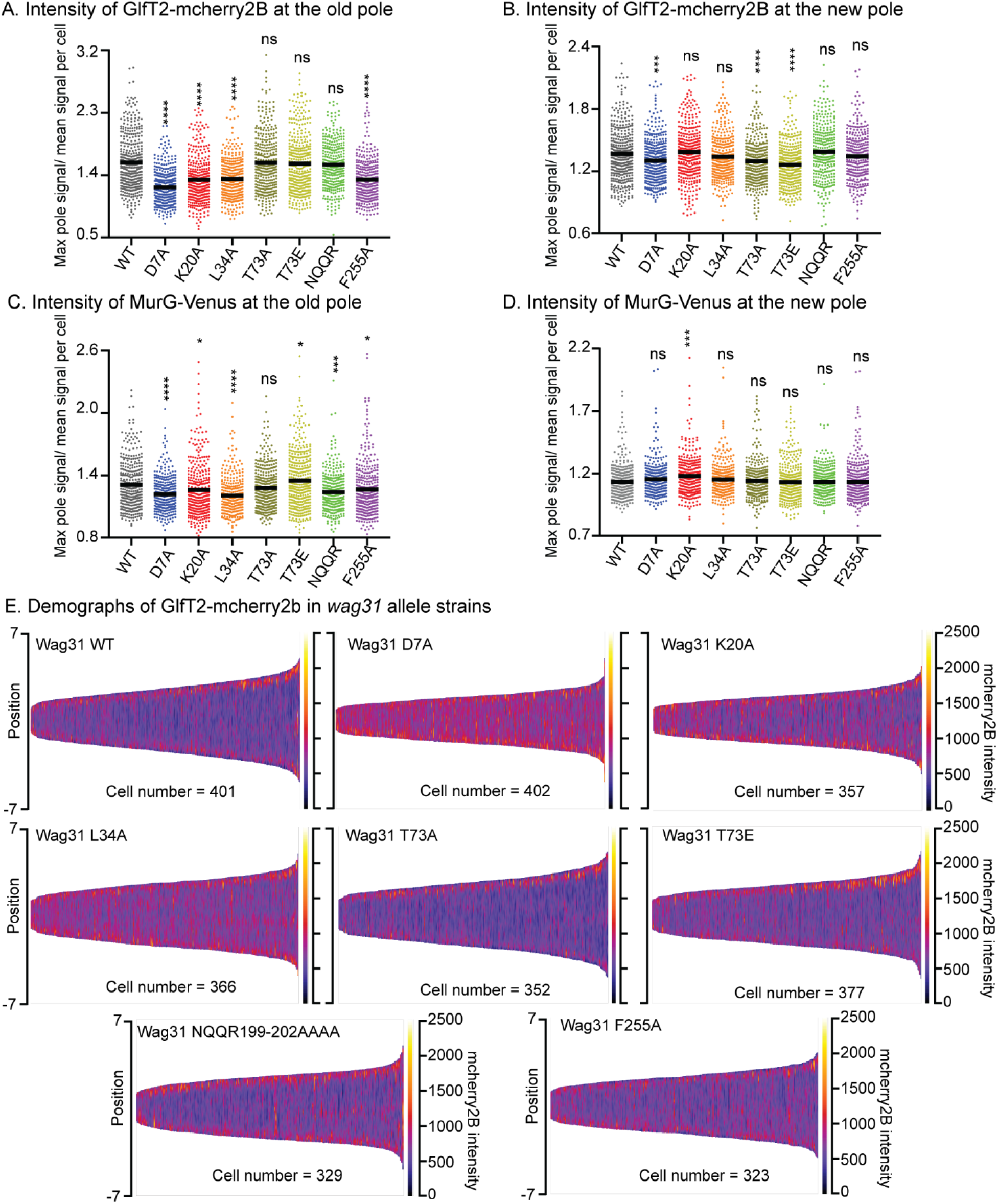
Localization of IMD proteins does not determine changes in polar growth. (A) and (B) Relative polar intensity of GlfT2-mcherry2b in the Wag31 WT and Wag31 mutants at the old pole and the new pole, respectively. (C) and (D) Relative polar intensity of MurG-Venus at the old pole and the new pole, respectively. (E) Demographs of GlfT2-mcherry2b intensity across the length of each cell (Y axis of each plot, with intensity indicated by color – scale to the right of each plot), arranged by cell length (along the X axis). At least 100 cells were analyzed for each of three biological replicate cultures. ns, P >0.05, *, P=< 0.05, ***, P= << 0.0005, ****, P=<0.0001. *P*-value is calculated by one-way ANOVA, Dunnett’s multiple comparisons test.

Across the mutants tested, polar elongation (Fig. 1HI) does not correlate either with GlfT2 or with MurG localization signals at the poles (Fig. 5). These data indicate that Wag31 may regulate the localization of IMD proteins, probably indirectly and largely at the old pole, but that this regulation does not directly control polar elongation.

## Discussion

Wag31’s molecular function in controlling cell wall metabolism has remained obscure. Homology with other systems predicts that Wag31 might recruit and regulate cell wall enzymes at the pole (8, 12). However, the hetero-oligomeric interactions that have been described for Wag31 – AccA3 (14, 39) and FtsI (40) - do not connect to Wag31’s role in restricting peptidoglycan metabolism to the poles. Additionally, these interactions may be indirect or only exist under certain stress conditions (40). Here, we sought to dissect Wag31’s function in regulating the cell wall (Fig. 6).

**Figure 6.**
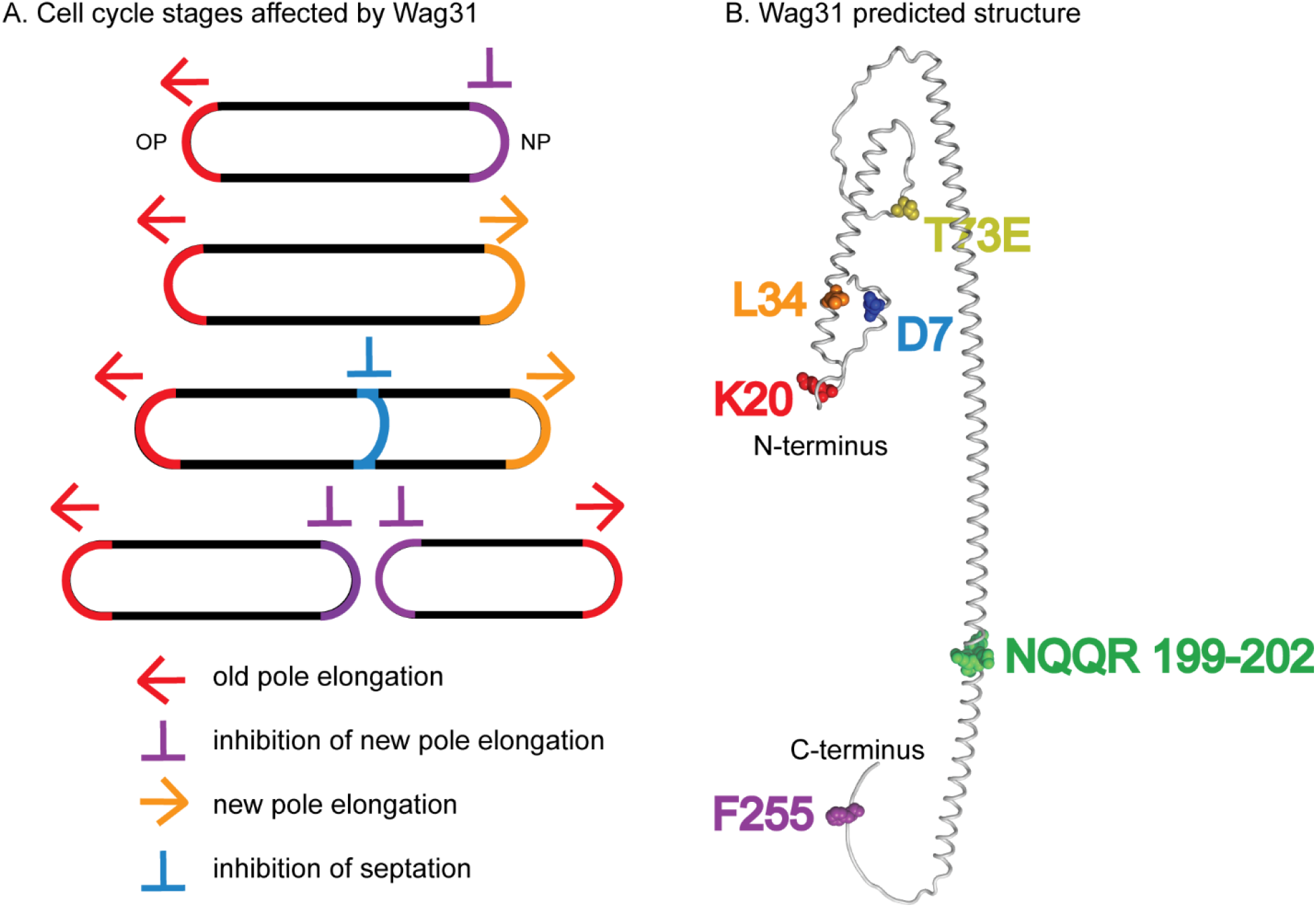
Models. (A) Multiple stages of the cell cycle affected by Wag31 are highlighted. (B) Alphafold2 predicted structure (Jumper *et al*., 2021) of a Wag31 monomer, with residues of interest highlighted.

Wag31 had previously been known to be essential for polar growth (6, 17). Here, we phenotypically profiled mutants of residues all along the length of Wag31 and found a variety of different phenotypes which indicate that Wag31 has distinct functions in polar elongation (Fig. 6AB). We found that the *wag31* K20A mutant is specifically more defective in old pole elongation while the *wag31* L34A mutant is specifically more defective in new pole elongation (Fig. 3). These results indicate that Wag31 is doing something different at the new and old poles; it may have a distinct homo-oligomeric conformations at the two poles, or may form different hetero-oligomeric regulatory interactions. Phosphorylation of Wag31 seems promote growth specifically at the old pole, as the phospho-mimetic *wag31* T73E mutant elongates more than the WT strain, while the phospho-ablative mutant elongates slightly less (Fig. 2H).

Our characterization of the *wag31* D7A mutant indicates that Wag31 is involved in downregulating septation. Wag31 had previously been shown to localize to the cell division site before septation (15), and overexpression of Wag31-GFP was shown to inhibit and mis-localize septa (13). Wag31 interacts with the septal PBP FtsI and regulates FtsI stability under oxidative stress through residues 46-48 (NSD) (40). We tested a *wag31* Δ46-48 mutant, as per (40) and found it was destabilized (Fig. S7). We also made a *wag31* NSD46-48AAA mutant, and found that it had normal cell length (Fig. S1), so those residues do not affect FtsI activity in exponential phase. The Wag31 D7 residue could possibly form a separate interaction with FtsI, or other septal factors (41). The phospho-site T73 on Wag31 may also have a role in regulating septation: we observed that the *wag31* T73A phospho-ablative mutant has slower elongation but longer cell length in log. phase (Fig. 2DFH), which suggests that it has delayed cell division. Delayed cell division of the *wag31* T73A is also apparent in stationary phase stress, as there are more cells with active peptidoglycan metabolism at the septum than in the WT strain (Fig. 3E).

We found that the *wag31* F255A mutant had increased polar elongation at the new pole (Fig. 1). This suggests that Wag31 has a role in inhibiting new pole elongation. The slower elongation at the new pole is partly controlled by membrane protein LamA (4). Our work suggests that either Wag31 works separately to inhibit new pole elongation, or that LamA may interact with Wag31 through F255 and interfere with Wag31’s promotion of new pole elongation.

Most of the mutants we characterized show a defect in GlfT2-mcherry2B and MurG-Venus localizations at the old pole, but not at the new pole. The localization of GlfT2 and MurG at the poles did not correlate with polar elongation defects (Fig. 3,5), indicating that polar localization of the IMD proteins is not the only determinant of polar growth. These data suggest that Wag31 is at best an indirect regulator of the IMD, and that regulation of the IMD is likely not Wag31’s essential function.

In this work, we define new roles for Wag31 in regulating both septation and the inhibition of the new pole, and identify residues that have dominant roles in Wag31’s different functions (Fig. 1, 6). We also describe how phosphorylation of Wag31 may regulate both old pole elongation and septation (Fig 2,3,6).

## Materials and Methods

### Bacterial strains and culture conditions

All *Msmeg* strains were grown in liquid culture in 7H9 (BD, Sparks, MD) medium supplemented with 0.2% glycerol, 0.05%Tween 80, and ADC (5g/L albumin, 2 g/L dextrose, 0.85 g/L NaCl, 0.003 g/L catalase). For most experiments, *Msmeg* strains were grown on LB Lennox plates. DH5a, TOP10, and XL1-Blue *E.coli* cells were used for cloning. For *E. coli*, antibiotics concentrations were: kanamycin – 50 <u>μ</u>g/ml; hygromycin - 100 <u>μ</u>g/ml; nourseothricin - 40 <u>μ</u>g/ml; Zeocin - 25 <u>μ</u>g/ml. For *Msmeg*, antibiotic concentrations were: kanamycin - 25 <u>μ</u>g/ml; hygromycin - 50 <u>μ</u>g/ml; nourseothricin - 20 <u>μ</u>g/ml; trimethoprim - 50 <u>μ</u>g/ml; Zeocin – 20 <u>μ</u>g/ml.

### Growth curves

Strains were grown to log. phase in 7H9 with appropriate antibiotics, then diluted to OD600 = 0.1 without antibiotics in a 96 well plate. A Synergy Neo2 linear Multi-Mode Reader was used to shake the plates continuously for 18 hours at 37°C, and read OD every 15 min. To find the doubling time for each strain, the raw data were analyzed with exponential growth equation model using GraphPad Prism (version 9.1.2).

### Strain construction

To build the allele replacement strains, first *wag31*_Msm_ under its native promotor was cloned into a kanamycin-marked L5 integrating vector and transformed into *Msmeg* mc^2^155. Then, *wag31* at its native locus was replaced with hygromycin resistance cassette by dsDNA recombineering (42). This cassette was oriented so the hygR promoter drives expression of downstream genes in the operon. Deletion of *wag31* from the genome was confirmed with PCR checks. *wag31*_Msm_ at the L5 site was swapped with *wag31*_Mtb_ alleles in a nourseothricin-marked L5-integrating vector using L5-phage site allelic exchange as described (32). The transformants were screened by antibiotic resistance in order to confirm allele swap. Vectors containing *wag31*_Mtb_ mutants were created using PCR stitching. All vectors were made using Gibson cloning (43). All strains, plasmids and primers used are listed in supplemental material part (Table S1A, S1B, S1C).

### Microscopy

All microscopy was performed on three biological replicate cultures of each strain using a Nikon Ti-2 widefield epifluorescence microscope with a Photometrics Prime 95B camera and a Plan Apo 100x, 1.45-numerical-aperture (NA) lens objective. Cells were taken from log. phase culture in 7H9 and immobilized on 1.5% agarose pads made with Hdb media. To detect GFPmut3 and Venus signal, a filter cube with a 470/40nm excitation filter, a 525/50nm emission filter and a 495nm dichroic mirror was used. To detect mCherry2B signal, a filter cube with a 560/40nm excitation filter, a 630/70nm emission filter and a 585nm dichroic mirror was used. To detect HADA, a filter cube with a 350/50nm excitation filter, a 460/50nm emission filter, and a 400nm dichroic mirror was used. Image analysis was performed using MicrobeJ (44) to make cell ROIs and extract fluorescence data from them. Fluorescence intensity data from MicrobeJ was further analyzed using bespoke MATLAB code (see supplement).

### Cell staining

Cultures were stained with 1 μg/mL HADA (R&D systems) for 15 minutes with rolling and incubation at 37°C. Stained cultures were then pelleted and washed in 7H9 before imaging. For the pulse-chase-pulse elongation test, cells after 15 minutes of HADA staining were resuspended into 7H9 for 1.5 hours of rolling and incubation at 37°C, then stained with 1 μg/mL NADA (R&D systems) for 2 minutes at room temperature, washed and resuspended in 7H9 before imaging.

## Acknowledgements

This work was supported by NIH grant R01AI148917 to CCB.

